# “Chronic Cervical Midline Contusion in Rats Disrupts Aerobic, Muscular, and Cardiovascular Function”

**DOI:** 10.1101/2025.08.24.671979

**Authors:** Z Yan, M Orellana, D Alarcon, RA German, AE Marcillo, T Pantelis, MS Nash, JD Guest, D McMillan, A Szeto, AJ Mendez, WW Muir, RL Hamlin, PD Ganzer

## Abstract

Cardiovascular dysfunction significantly contributes to morbidity and mortality following cervical spinal cord injury (SCI). Unfortunately, only a limited number of preclinical models have been developed for investigating cardiovascular dysfunction following cervical SCI. Furthermore, the broader consequences of cervical SCI on aerobic capacity and muscle endurance during physiological stress testing also remains understudied preclinically. Therefore, in this study we assessed potential deficits across multiple physiological systems in a rat model of cervical SCI using a battery of stress tests. Female Sprague-Dawley rats (n = 20) received either a C8 midline contusion (cSCI) or laminectomy alone as a control (LAM). Exercise stress testing was conducted to evaluate cardiorespiratory fitness and recovery using a metabolic treadmill, or forelimb fitness using the isometric pull task. Orthostatic stress testing and pharmacological stress testing were also performed to more directly challenge the cardiovascular system. Our findings demonstrate a decline in aerobic fitness in cSCI rats, as evidenced by dysregulated excess post-exercise oxygen consumption. cSCI rats also exhibited impaired muscle endurance compared to LAM. During orthostatic stress testing, 70% of cSCI rats experienced an approximately 25 mmHg decrease in systolic blood pressure and 20 mmHg decrease in diastolic blood pressure, in addition to modest but significant decreases in heart rate, myocardial contractility index, stroke volume index, and cardiac output index. During dobutamine infusion, cardiac output index and stroke volume index were significantly reduced following cSCI compared to LAM. Overall, these stress testing results suggest that preclinical cervical SCI in rats can lead to, and therefore model, clinically relevant impairments in cardiopulmonary exercise performance, muscle endurance, and cardiovascular function.

## Introduction

Cervical spinal cord injury (SCI) is the most common cause of SCI and can lead to debilitating deficits across multiple physiological systems (1–4). Many human studies have demonstrated impaired aerobic function, decreased muscle endurance, orthostatic dysregulation, and cardiovascular dysfunction following cervical SCI (1,5–7). These significant impairments can occur following both complete, and interestingly, incomplete SCI (1,2,5,7). Regarding preclinical models, the evaluation of aerobic and cardiovascular changes following SCI appears to have progressed at a slower pace compared to the evaluation of other systems (such as the well-studied sensorimotor system). Overall, there is a major need to develop clinically relevant models of aerobic, muscular, and cardiovascular dysfunction following cervical SCI.

Cervical SCI impacts several physiological systems, including cardiovascular control, via damage to the descending sympathetic outflow control pathways (3,8–10). The majority of supraspinal sympathetic outflow control fibers originate in the rostral ventrolateral medulla (2,11–13), in addition to other supraspinal areas such as the caudal raphe nucleus and the A5 cell group (2,12,14–16). This descending control system is complex and generally projects via the dorsolateral funiculi bilaterally and diffusely to the preganglionic sympathetic neurons located in the intermediolateral cell column (roughly spanning spinal levels T1-L2 (2,3,8)). Damage to these descending pathways can significantly impact responses to physiological challenges or other stressors in part due to impaired cardiovascular function.

Cardiovascular function can be assessed using aerobic or resistance exercise stress testing (6,17–19). Aerobic function is commonly measured by recording inspired and expired gases during maximal exercise testing. This enables the measurement of peak volume of oxygen consumption (e.g., VO_2 peak_), a gold standard for assessing aerobic capacity (19–23), as well as excess post-exercise oxygen consumption (EPOC) during recovery, a reliable indicator of aerobic fitness (24–30). Resistance exercise stress testing can also be employed to challenge the musculoskeletal and cardiovascular systems. Furthermore, there are other alternative stress tests that more directly challenge the cardiovascular system (e.g., in the absence of significant musculoskeletal contributions). Orthostatic stress (i.e., the stress of assuming an upright position) is widely used clinically and preclinically and commonly involves the use of a tilt table (6,31–33). Lastly, pharmacological stress testing can also be used to challenge cardiovascular function (e.g., using a dobutamine infusion) (34–38).

In the current study, we hypothesized that chronic incomplete cervical SCI in rats would disrupt aerobic function, muscle endurance, and multiple facets of cardiovascular control. Approximately 3-4 months after injury, rats with a cervical SCI (cSCI) or a laminectomy (LAM) were exposed to: 1) aerobic exercise stress testing using a metabolic treadmill, 2) resistance exercise stress testing using the isometric pull task, 3) orthostatic stress testing using a tilt table, and 4) pharmacological stress testing using dobutamine infusions. Our overall results characterize a clinically relevant model of incomplete cervical SCI in rats that can be used to investigate cardiovascular dysfunction and the effect of therapeutic interventions.

## Methods

### Overview

All procedures were approved by the Institutional Animal Care and Use Committee of the University of Miami (Miami, FL). Adult female Sprague Dawley rats (∼300 g; n = 20) used in this study were housed one per cage (12 hr light/dark cycle; *ad libitum* access to food and water). The general aims of the study were to assess the effects of chronic cervical midline contusion on aerobic function, forelimb fitness, and cardiovascular function. An overview of the study’s techniques and experimental timelines are shown in Fig. 1. All rats were first trained to proficiency on the metabolic treadmill and isometric pull tasks prior to surgery. Rats were then randomized into balanced groups receiving either a cervical contusion at spinal level C8 (cSCI group) or laminectomy as a control (LAM group). Rats were then left sedentary in their home cage until further testing. Exercise stress testing, including resistance exercise using the isometric pull task and aerobic exercise using the metabolic treadmill, was performed at Week 13 and Week 14 post-surgery, respectively. Orthostatic stress testing and pharmacological stress testing occurred during Week 16 post-surgery. All data analyses were performed in MATLAB or GraphPad Prism.

**Fig. 1.**
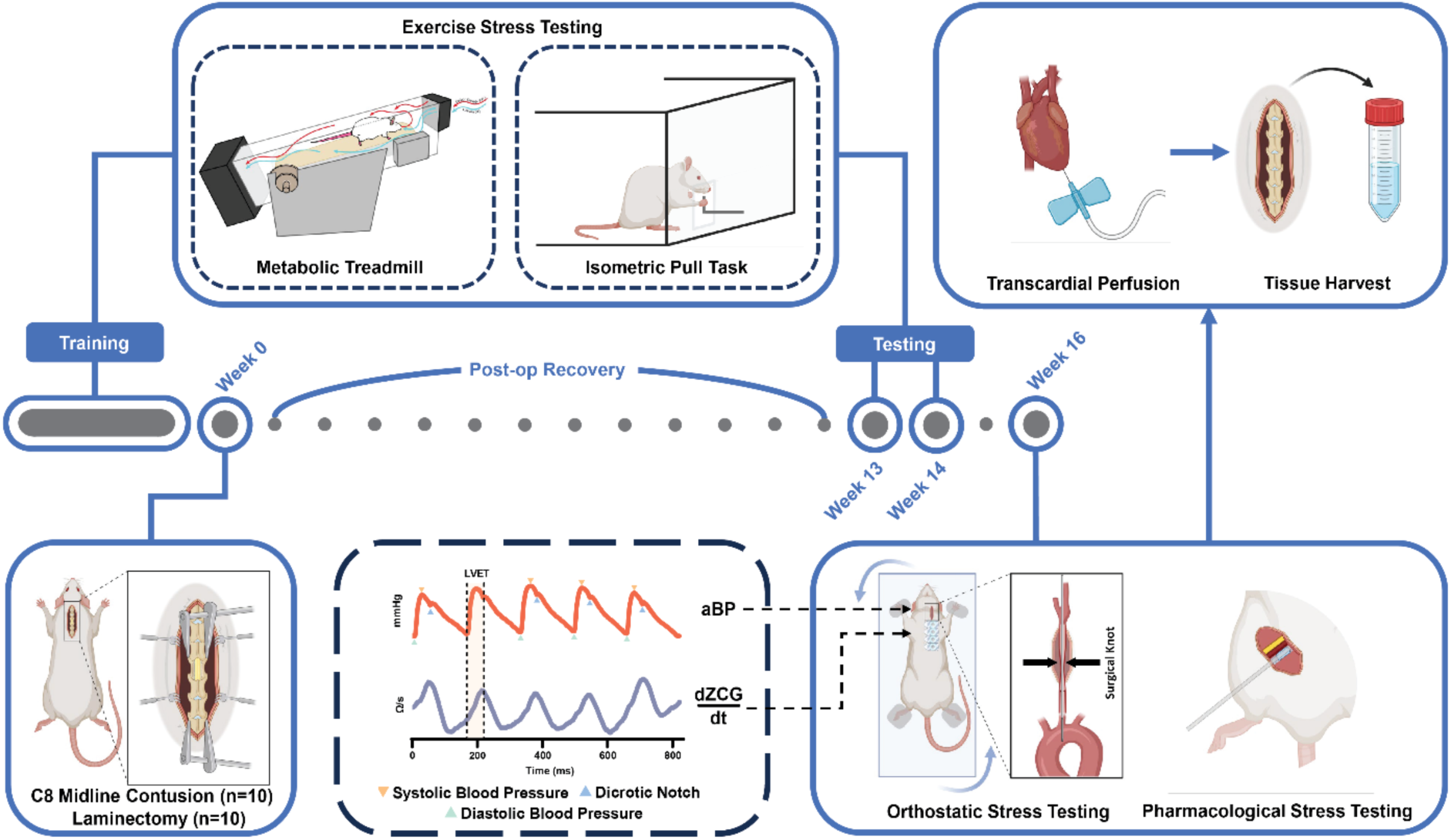
Study Overview: Methods and Timeline. Cartoon schematics of the exercise stress testing paradigms (**top left**), surgery (**bottom left**), the orthostatic and pharmacological stress testing paradigms (**bottom right**), tissue collection procedures (**top right**), and study timeline (**middle**). (LVET: left ventricular ejection time; aBP: arterial blood pressure; mmHg: millimeters of mercury;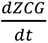: the first derivative of the impedance cardiography (ZCG) signal; Ω: ohms). Cartoon schematics made in part with BioRender.

### C8 Midline SCI or Laminectomy Surgery

Ten rats received a cervical midline contusive SCI using previously reported methods (39), whereas 10 control rats received a laminectomy alone (Fig. 1, bottom left). Rats were first anesthetized with isoflurane (1.5 - 2% vol.). The back of the animal was then shaved and cleaned. Incisions were made through the dorsal muscles to expose the cervical spinal column. Partial laminectomies were performed to expose spinal level C8. The spinal column was then stabilized, and a midline contusion was produced using a force of 250 kilodynes and zero dwell time (Infinite Horizon Impact Device, Precision Systems and Instrumentation; impactor tip diameter = 2.5 mm). The muscle overlying the exposed vertebra was then closed in layers and the incised skin closed using surgical staples. All rats received post-operative fluids (lactated ringers electrolyte solution; s.c.), antibiotics (enrofloxacin, 10 mg/kg; s.c.), and analgesics (buprenorphine extended release, 1 mg/kg; s.c.) immediately after surgery. Fluid and antibiotic administration continued for 3 days post-operatively. Animal bladders were expressed as needed until bladder function returned to normal. One rat in the LAM group was sacrificed due to post-operative complications.

### Metabolic Treadmill Exercise Stress Testing

A motorized metabolic treadmill (0184-006M; Columbus Instruments; Columbus, Ohio) was used to assess aerobic function during Week 14 post-surgery (Fig. 1, top left). The treadmill inclination was maintained at 5 degrees and gas sensors were calibrated prior to each testing session. Treadmill acclimatization and VO_2 peak_ testing protocols were adapted from Koch and Britton (19), a widely used set of methods in the preclinical aerobic exercise performance literature (40–43). Briefly, prior to surgery, all rats were gradually acclimatized to the motorized metabolic treadmill, by ramping up treadmill speed, running duration, and shock grid intensity across a 5-day period. VO_2 peak_ testing was then performed twice, separated by one day. A VO_2 peak_ testing session began with a 300 s baseline recording at a treadmill speed of 0 m/min. The treadmill then increased to 3 m/min, followed by 10 m/min, with each speed held for 120 s. Subsequently, the treadmill accelerated incrementally by 2 m/min intervals, with each speed maintained for 120 s. The shock grid intensity was maintained at 1.6 mA for all VO_2 peak_ testing sessions to motivate running. After the rats remained on the shock grid for at least 7 seconds, indicating exhaustion and the end of VO_2 peak_ testing, the treadmill speed was reduced to 3 m/min and held for a 300 s cool-down post-exercise period. Excess post-exercise oxygen consumption (EPOC) was assessed based on the slope of VO_2_ change during the first minute of the cool-down period, describing the rapid component of EPOC (28–30). Similar to previous studies (19), for the VO_2 peak_ testing day with the best performance, we report the peak VO_2_ achieved at the end of exercise testing (i.e., VO_2 peak_ (mL/[kg/hr])), EPOC (mL/[kg/hr^2^]), time to exhaustion (s), and total distance ran (m). These metrics were compared across cSCI and LAM rats to assess aerobic capacity and fitness.

### Isometric Pull Task

The isometric pull task was performed to evaluate multiple measures of forelimb fitness (Fig. 1, top left). We specifically utilized the isometric pull task here to also examine muscle endurance, similar to previous muscle endurance studies using maximal voluntary contraction resistance exercise, requiring near maximal force exertion and high trial counts over time (44–47). A given animal performed an average of 515 ± 65 isometric pulls total during a given 30-minute testing session, allowing for sufficient peak force decay and subsequent quantification of muscle endurance. All rats were trained to proficiency using a 120 g forelimb pull force success threshold, as described in our previous studies (39). At Week 13, forelimb fitness metrics (number of trials, peak force (g), and force velocity_max_ (g/ms), as well as forelimb muscle endurance measures (Δ number of trials, Δ peak force (g), and Δ force velocity_max_ (g/ms) between 1^st^ and 4^th^ quartile of the task session) were assessed using a challenging (120 g) or a submaximal (40 g) forelimb pull force success threshold on separate testing days.

### Cardiovascular Sensor Instrumentation

Approximately 2 weeks after VO_2 peak_ testing, rats were next instrumented with cardiovascular sensors for orthostatic stress testing and pharmacological stress testing (Fig. 1, bottom right). Rats were first anesthetized using urethane (1.3-1.5 g/kg; i.p.). Supplemental anesthesia was administered if needed and the surgical plane was kept at stage III-3 (48). Rats were then transferred onto a heating pad (Stoelting Co, IL) and the rectal temperature was maintained at ∼36-38 ℃ for the duration of the procedure. The neck, the thorax, and the chest region were shaved. Blunt dissection was next performed over the left cervical carotid area to expose the left carotid artery. A calibrated pressure catheter (model SPR-407, Millar, Inc., TX) was inserted into the left common carotid artery, advanced approximately 2 cm deep into the aortic arch, and finally secured with suture allowing for the measurement of arterial blood pressure (aBP).

Two pairs of impedance cardiography (ZCG) electrodes (EL501, BIOPAC systems Inc., CA) were next placed on the left neck and chest region (49,50). Electrode gel (BIOPAC systems Inc., CA) was applied to the electrodes to ensure optimal electrical conductivity and recording quality. ZCG provides a non-invasive estimation of cardiovascular metrics, including stroke volume (SV) and SV index (49–53).

### Orthostatic Stress Testing

Rats were next transferred to a tilt table and placed supine. To maintain the head and body position during tilting, all four limbs were then affixed onto the heating pad with surgical tape (3 M, Maplewood, MN). Cardiovascular data was monitored during supine, ∼90° head-down tilt (or H-DT, allowing for baroreceptor loading), or ∼90° head-up tilt positions (or H-UT, allowing for baroreceptor unloading). An orthostatic stress testing session (Fig. 1, bottom right) started at the baseline supine position, followed by two H-UTs and one H-DT in a random sequence. Rats returned to the baseline supine position between each tilting epoch. Cardiovascular data recording occurred continuously, and each position lasted approximately 3.5 minutes.

### Pharmacological Stress Testing

Following orthostatic stress testing, the right femoral vein was dissected and cannulated allowing for intravenous administration of dobutamine (DOB, 2 µg/kg/min and 10 µg/kg/min; Fig. 1, bottom right), similar to our previous work (38). We recorded cardiovascular data during a 30 s baseline period prior to a given infusion. Each infusion period lasted 5 min, followed by a 5 min post-infusion recovery period. Between each infusion, cardiovascular states returned to baseline levels. Responses to DOB infusion were assessed from the baseline period to the end of infusion.

### Cardiovascular Signal Processing

Simultaneous recordings of aBP and ZCG were sampled at 1 kHz during the orthostatic and pharmacological stress testing procedures (Fig. 1, bottom middle). The recordings were converted into continuous signals for offline analysis. 60 Hz noise was removed from all recorded signals using a notch filter. The ZCG signal was further processed to derive the following cardiovascular measures: myocardial contractility index (C index; [Ω/s]/m^2^)), stroke volume index (SV index; [mL/beat]/m^2^), and cardiac output index (CO index; [L/min]/m^2^), similar to previous studies (49,54–57). The distance between two inner electrodes (L; cm) was measured to calculate the volume of electrically participating tissue (VEPT; mL) using the 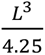 ratio (49). Breath motion artifacts were automatically detected in MATLAB. 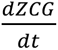 was next computed and data points lying 0.2 s before and after a given breath motion artifact were rejected to further optimize overall ZCG signal analysis. 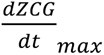 (Ω/s) is defined as the maximal change from the minima to the peak of the 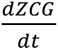 signal during systole (56,58,59). Stroke volume (SV; mL/beat) was calculated using the following formula used in previous work (49,52,53): 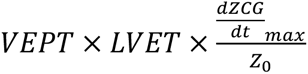 (where LVET (s) is the left ventricular ejection time, equal to the interval from the beginning of systole to the dicrotic notch in the aortic aBP waveform from the corresponding beat; *Z_o_* (Ω) is the base impedance of the ZCG signal). Cardiac output (CO; L/min) was calculated by multiplying heart rate (HR; bpm) and SV. Vallois formula was used to estimate body surface area (BSA; m^2^) using the body weight and nose-to-tail length of rats (49,60). C index, SV index, and CO index were calculated by 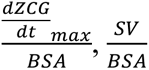, and 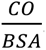. HR was estimated in MATLAB using detected systolic peaks from the aBP signal (Fig. 1, bottom middle). The custom MATLAB program also allowed for the detection of systolic blood pressure (SBP, mmHg) and diastolic blood pressure (DBP, mmHg). Therefore, pulse pressure (PP, mmHg; *SBP* − *DBP*) and mean arterial pressure 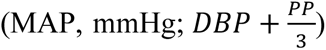 were also calculated. All data were binned in one second intervals and missing values were interpolated if needed using a shape-preserving piecewise cubic method. Signal changes (Δ) were calculated by subtracting mean baseline values.

### Blood Volume Estimation

Hematocrit (%) has been used to estimate blood volume, calculated by measuring the length of the packed red cells divided by the length of whole blood volume in a centrifuged microhematocrit tube (61–65). At Week 16, blood samples were collected into microhematocrit tubes with heparin (VWR, PA) and centrifuged for 5 mins at 7200 rpm (Clay Adams Readacrit Centrifuge; Becton Dickson, NJ). The length of the packed red cells and whole blood volume was then measured to calculate hematocrit (%). A higher hematocrit (%) is indicative of a lower blood volume (62–65).

### Transcardial Perfusion and Tissue Collection

Rats were next euthanized using a urethane overdose. A butterfly needle was inserted into left ventricle and the rat was transcardially perfused using 0.01 M phosphate buffered saline (PBS) (stored at 4 ℃ before use) at a rate of 10 mL/min for 10 mins (∼100 mL). Once the perfusion began, the right atrium was cut to open the circulatory system. After the perfusion, the spinal cord was dissected and fixed in 4% paraformaldehyde (PFA) at 4 ℃ for 7 days. The harvested spinal cord lengths were 10 mm (5 mm rostral and caudal from the C8 SCI epicenter). All samples were stored in 0.01 M PBS with 0.03% sodium-azide (NaN_3_) at 4 ℃ until tissue clearing, staining, and imaging.

### Spinal Cord Tissue Processing and cSCI Lesion Assessment

Spinal cord tissue was cleared and stained using a modified immunolabeling-enabled three-dimensional imaging of solvent-cleared organs (iDISCO) protocol (66). Spinal cord tissues were first dehydrated in a graded methanol series (20%, 40%, 60%, 80% of methanol in distilled water) for 1 hour each. Following dehydration, samples were transferred into 100% methanol twice for 1 hour each and chilled at 4 ℃. Samples were then placed into a solution of 66% dichloromethane (DCM) and 33% methanol overnight with agitation at room temperature. On the second day, the samples were washed twice with 100% methanol, each for 1 hour at room temperature. The samples were next bleached by 5% H_2_O_2_ in methanol (vol/vol) overnight at 4 ℃. After bleaching, the samples were rehydrated in a graded methanol series, each for 1 hour. This was followed by a wash at 0.01 M PBS and two washes in a solution of 0.01 M PBS with 0.2% TritonX-100, each for 1 hour at room temperature. To enable immunolabeling, the samples were incubated in permeabilization solution (0.01 M PBS with 0.2% Triton X-100, 20% dimethyl sulfoxide (DMSO), and 0.3 M glycine) and blocking solution (0.01 M PBS with 0.2% Triton X-100, 10% DMSO, and 5% donkey serum) at 37 ℃, each for 2 days. Next, samples were cut into 1 mm pieces and incubated in anti-myelin primary antibody (Biolegend, ab2814552, 1:200) diluted in 5% DMSO, 3% donkey serum, and 0.01M PBS with 0.2% Tween-20 and 10 mg/mL heparin (PTwH) for 1 week at 37 °C with agitation, and then washed several times in PTwH. Immunolabeled samples were then dehydrated in a graded methanol series, followed by incubation in 66% DCM and 33% methanol for 3 hours with agitation and in 100% DCM twice, each for 15 minutes at room temperature to wash out methanol. Finally, the samples were cleared in diBenzyl ether (DBE) until imaging.

The samples were finally imaged using a Andor Dragonfly 200 confocal microscope (Oxford Instruments, United Kingdom) at 10x magnification. Images were loaded into a custom MATLAB application to quantify lesion extent. We report the percentage of spared white and grey matter 5 mm rostral and 5 mm caudal from the lesion epicenter at 500 µm intervals.

### Statistical Analyses

All statistical analyses were performed using GraphPad Prism 10 (GraphPad Software, San Diego, CA). Continuous variables are presented as mean ± standard error of the mean (S.E.M.). The significance level (alpha) was set at 0.05. The normality of all data was first examined, and non-parametric tests were used if needed (e.g., Mann-Whitney tests). To determine aerobic function and forelimb fitness pre-surgery, two-tail unpaired t-tests or Mann-Whitney tests were used. A two-way ANOVA (group × threshold) with multiple comparisons using uncorrected Fisher’s Least Significant Difference (LSD) was used to examine forelimb fitness metrics and correlates of muscle endurance at Week 13. To assess aerobic function metrics at Week 14, two-tail unpaired t-tests or Manny Whitney tests were used. Pearson’s correlation was performed to assess the relationship between VO_2 peak_ and EPOC. Two-tailed tests were performed on all baseline cardiovascular data. Two-way repeated-measures ANOVA (group × time) with Bonferroni’s multiple comparison were used to analyze the effect of orthostatic stress testing on cardiovascular states across time. The area under the curve (AUC, a.u.) was calculated for cardiovascular data during pharmacological stress testing. Parametric two-way ANOVAs (group × dose) with multiple comparisons using uncorrected Fisher’s LSD were used to analyze the effect of DOB on cardiovascular function.

## Results

### Study Overview

Aerobic function, forelimb fitness, and cardiovascular function were assessed in two groups of rats (cSCI and LAM; study overview: Fig. 1). All rats were trained to perform two different exercise stress tests prior to cSCI or LAM surgery, followed by subsequent testing again at Week 13 and Week 14. Orthostatic and pharmacological stress testing were performed during an anesthetized terminal procedure with cardiovascular sensor instrumentation during Week 16.

### cSCI Lesion Assessment in Cleared Spinal Tissue

SCI at spinal level C8 was performed in order to damage functionally relevant pathways, including the descending sympathoexcitatory pathways innervating the interomediolateral column (IML) contributing to cardiovascular control (Fig. 2A; (3)). At the end of the 16 week study, spinal tissues were cleared and stained for myelin using the iDISCO protocol (66), allowing for spinal cord lesion assessment. At the cSCI lesion core (Fig. 2B, right), there was approximately 10% spared gray matter and 27% spared white matter (Fig. 2C).

**Fig. 2.**
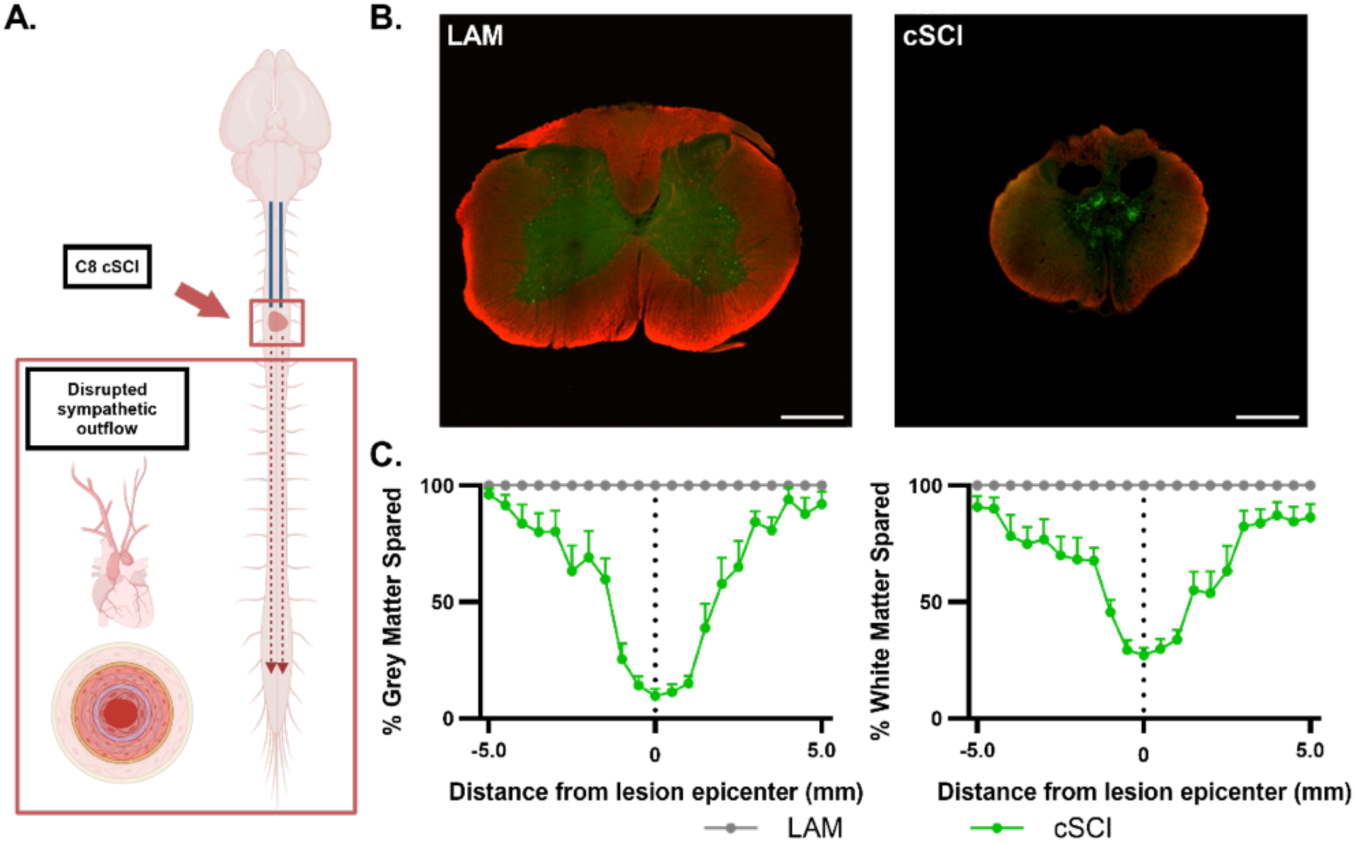
cSCI Lesion Analysis. **A**. Cartoon schematic of the rat neuraxis and the C8 cSCI disrupting multiple descending pathways, including sympathetic control circuits. **B**. Representative images of cleared tissue stained for myelin using the iDISCO protocol from a representative LAM rat and a representative cSCI rat (at spinal level C8). Scale bar = 400 µm. **C**. Rostrocaudal distribution of spared gray matter and spared white matter across groups (rostral = + mm; caudal = - mm; mean ± S.E.M.). Cartoon schematic made in part with BioRender (panel **A**).

### cSCI Impairs Aerobic Fitness and Forelimb Muscle Endurance

Aerobic function was assessed during aerobic exercise stress testing using a metabolic treadmill. We assessed the hypothesis that cSCI would significantly decrease measures of aerobic capacity and aerobic fitness. Interestingly, cSCI did not significantly impair VO_2 peak_ compared to LAM (Fig. 3A, unpaired Mann-Whitney test, Mann-Whitney U = 27, *p* = 0.274). Overall, LAM rats achieved a similar VO_2 peak_ level when compared to weight- and age-matched Sprague Dawley rats from previous studies (67,68). In contrast, EPOC (Fig. 3B, unpaired Student’s t-test, t(16) = 2.59, *p* = 0.0198), total time to exhaustion (Fig. 3C, unpaired Student’s t-test, t(16) = 2.59, *p* = 0. 0195), and total distance ran (Fig. 3D, unpaired Student’s t-test, t(16) = 2.30, *p* = 0.0354) were significantly impaired in cSCI rats. In addition, VO_2 peak_ was negatively correlated with EPOC, indicating that animals with a lower aerobic capacity also exhibited dysregulated post-exercise recovery (R = −0.683, *p* < 0.01). Basso, Beattie, and Bresnahan (BBB) scores demonstrated that cSCI rats exhibited consistent weight-supported stepping needed for performing aerobic exercise stress testing (Fig. S1). These results indicate that correlates of aerobic function are moderately impaired following cSCI in rats.

**Fig. 3.**
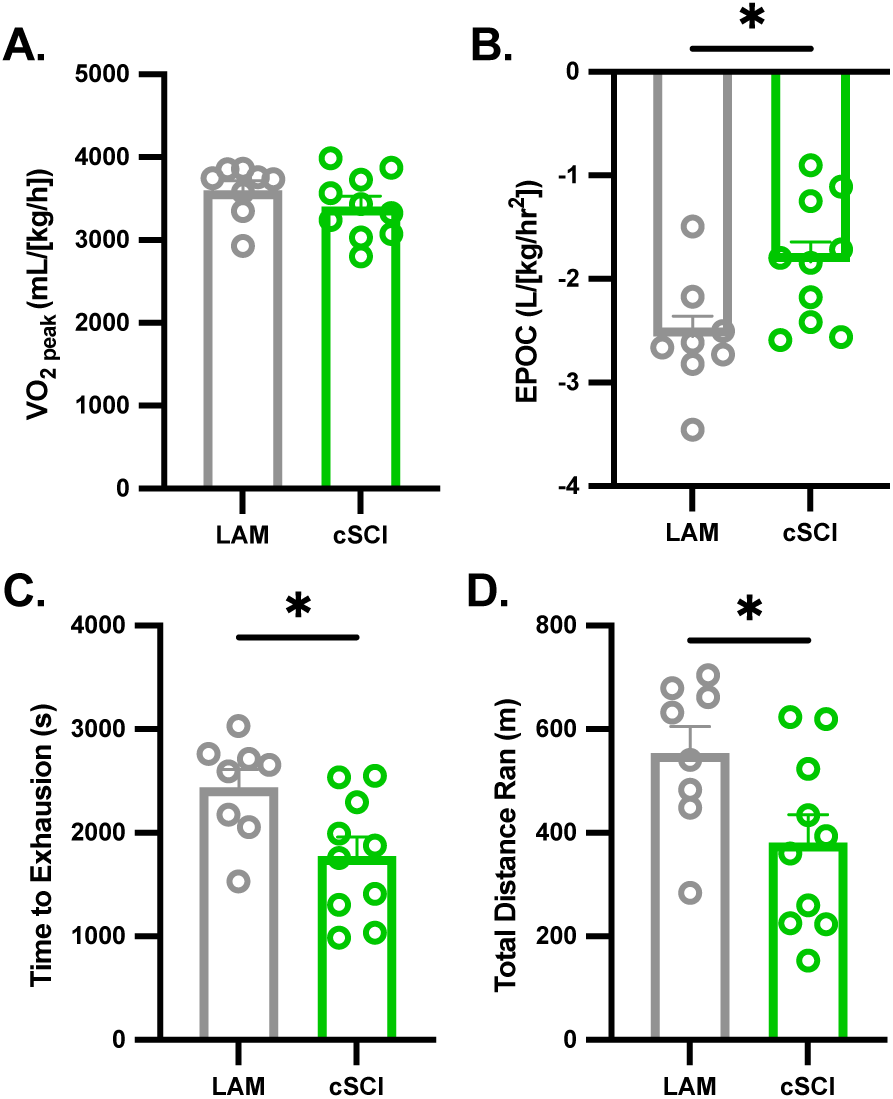
cSCI Significantly Impacts Aerobic Fitness and Correlates of Exercise Tolerance During Metabolic Treadmill Testing. cSCI rats demonstrated a trend towards a decrease in VO_2 peak_ (**A**), as well as significant impairments in excess post-exercise oxygen consumption (EPOC; **B**), time to exhaustion (**C**), and total distance ran (**D**) (\**p* < 0.05; mean ± S.E.M.).

**Fig. S1.**
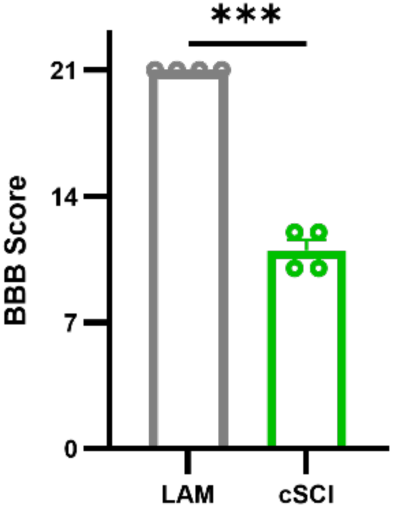
Basso, Beattie, Bresnahan (BBB) Scores Demonstrating Frequent to Consistent Weight-supported Stepping in A Subset of cSCI rats (\*\*\**p* < 0.001; mean ± S.E.M.).

Forelimb strength was assessed during resistance exercise stress testing using the isometric pull task. Due to the level of the injury at spinal level C8, we hypothesized that cSCI would have minimal effects on forelimb strength metrics (e.g., peak force), but would significantly affect forelimb muscle endurance (via damage to the descending sympathetic outflow pathways innervating the thoracolumbar IML and possible effects on cardiovascular function). Rats were first trained to proficiency on the isometric pull task (39). There were no differences in any isometric pull task metric pre-surgery across groups (Unpaired Student’s t-test, number of trials: t(17) = 1.81, *p* = 0.0886; unpaired Mann-Whitney test, peak force: Mann-Whitney U = 40, *p* = 0.720; unpaired Welch’s t-test, force velocity_max_: t(10.2) = 1.17, *p* = 0.126). At Week 13, rats were tested using a challenging maximal (120 g) or a submaximal (40 g) forelimb pull force success threshold, similar to previous studies using maximum voluntary contraction thresholds for assessing muscle fitness (44,45). There were no significant differences in the number of trials at either pull force success threshold (Fig. 4A; two-way ANOVA, F[1, 28] = 0.0487, *p* = 0.827). cSCI rats exhibited a moderate but significant deficit in force velocity_max_ at both pull force success thresholds compared to LAM (Fig. 4B; two-way ANOVA, Main effect; F[1, 28] = 9.096, *p* = 0.0054). Importantly, there were no significant differences in peak force across groups at either pull force success threshold (Fig. 4C; two-way ANOVA, F[1, 28] = 0.0840, *p* = 0.774).

**Fig. 4.**
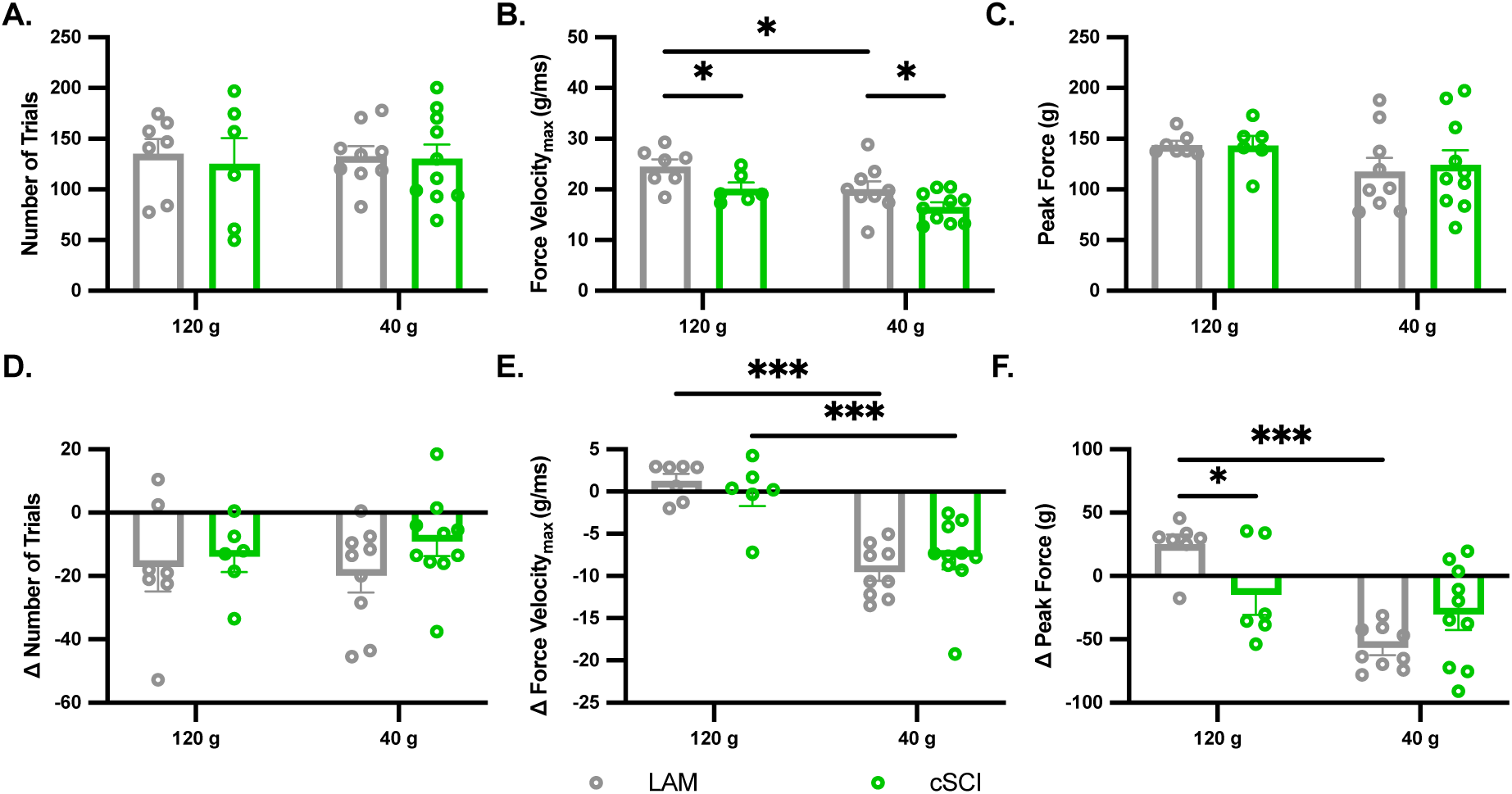
cSCI Significantly Impairs Muscle Endurance During the Isometric Pull Task. **A**. cSCI and LAM rats demonstrated a similar number of trials across pull success testing thresholds. **B**. cSCI rats exhibited significant decreases in force velocity_max_ at both pull success testing thresholds compared to LAM. **C**. There were no differences in peak force across groups at either pull success testing threshold. **D**. There were no differences in Δ number of trials. Δ Force velocity_max_ decayed significantly more at the lower submaximal pull success testing threshold (40 g) within groups. **F**. Importantly, when challenged at a maximal force threshold (120 g), cSCI rats showed a significant deficit in Δ peak force decay, indicative of impaired muscle endurance (\**p* < 0.05; \*\*\**p* < 0.01; mean ± S.E.M.).

Forelimb muscle endurance was next measured by quantifying the difference (Δ) in peak force from the beginning to the end of a given 30-minute assessment period, similar to previous studies on muscle endurance (69,70). We hypothesized that cSCI would significantly decrease forelimb muscle endurance. Muscle endurance metrics did not differ between groups pre-surgery (unpaired Welch’s t-test, Δ number of trials, *p* = 0.521; unpaired Welch’s t-test, Δ peak force, *p* = 0.337; unpaired Mann-Whitney t-test, Δ force velocity_max_, *p* = 0.905). At Week 13, there was no difference in Δ number of trials or Δ force velocity_max_ across groups at either pull force success threshold across groups (Fig. 4D; two-way ANOVA, F[1, 28] = 0.430, *p* = 0.517; Fig. 4E; two-way ANOVA, F[1, 28] = 0.302, *p* = 0.587). As expected, both cSCI and LAM rats pulled significantly slower when exposed to the less challenging submaximal forelimb pull force success threshold of 40 g (Fig. 4E; two-way ANOVA, Main effect; F[(1, 28) = 8.65, *p* = 0.0065). Importantly, cSCI rats demonstrated correlates of impaired forelimb muscle endurance, exhibiting a significantly larger decrease in Δ peak force compared to LAM at the challenging maximal pull force success threshold of 120 g (Fig. 4F; two-way ANOVA, F[(1, 28) = 4.259, p = 0.0484). Overall, these results support the hypothesis that multiple facets of muscle fitness are impaired following cSCI.

### cSCI Leads to Significant Cardiovascular Dysfunction and Orthostatic Hypotension

Cardiovascular control was next measured during orthostatic stress testing using a tilt table (Fig. 1, bottom right). Systolic blood pressure was significantly lower in cSCI rats at baseline prior to orthostatic stress testing, while other cardiovascular measures were not different across groups (Table 1). Importantly, there was no significant difference in estimated total blood volume across groups (62–65) (Fig. S2; Unpaired t-test, t(10) = 1.62, *p* = 0.135). Orthostatic stress testing was next performed, where body position changes enabled loading (head-down tilt or H-DT) or unloading (head-up tilt or H-UT) of the arterial baroreceptors. We hypothesized that cSCI would significantly impair sympathoexcitatory capacity and cardiovascular function during orthostatic stress (i.e., H-UT).

**Table 1.** Overview of Baseline Measures for Orthostatic Stress Testing. (MAP: mean arterial pressure; SBP: systolic blood pressure; DBP: diastolic blood pressure; PP: pulse pressure; HR: heart rate; LVET: left ventricular ejection time; C index: myocardial contractility index; SV index: stroke volume index; CO index: cardiac output index; mean ± S.E.M.; Mann-Whitney t-test: ^†^ *p*<0.05).

| Measures | LAM | cSCI | *p* |
| --- | --- | --- | --- |
| MAP (mmHg) | 93.6 ± 5.52 | 80.6 ± 5.07 | 0.0947 |
| SBP (mmHg) | 121 ± 4.82 | 106 ± 4.69 | 0.0435<sup>†</sup> |
| DBP (mmHg) | 74.7 ± 5.90 | 67.8 ± 5.35 | 0.153 |
| PP (mmHg) | 41.5 ± 1.48 | 37.8 ± 1.73 | 0.125 |
| HR (bpm) | 347 ± 13.4 | 367 ± 17.5 | 0.381 |
| LVET (s) | 0.0514 ± 0.00255 | 0.0492 ± 0.00187 | 0.483 |
| C index ([$\Omega$/s]/m<sup>2</sup>) | 2.85 ± 0.722 | 3.08 ± 0.533 | 0.803 |
| SV index ([mL/beat]/m<sup>2</sup>) | 0.290 ± 0.0800 | 0.279 ± 0.0516 | 0.911 |
| CO index ([L/min]/m<sup>2</sup>) | 0.103 ± 0.0293 | 0.104 ± 0.0187 | 0.977 |

**Fig. S2.**
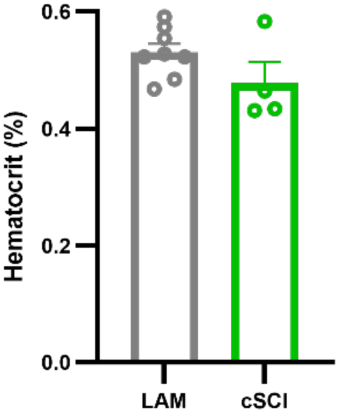
cSCI Does Not Affect The Estimated Total Blood Volume Using The Hematocrit Percentage Method (mean ± S.E.M.).

During orthostatic stress (H-UT), cSCI rats demonstrated a significant inability to raise heart rate (HR) by the end of the tilt period compared to LAM (Fig. 5A; two-way repeated ANOVA, Interaction; F[6, 102] = 4.66, *p* < 0.001). H-UT also led to a significant drop in mean arterial pressure (MAP) for cSCI (∼20 mmHg by the end of the H-UT period), ∼4-5 fold more compared to LAM (Fig. 5B; two-way repeated ANOVA, F[6, 102] = 9.70, *p* < 0.001). This was also accompanied by significant decreases in systolic blood pressure (SBP) and diastolic blood pressure (DBP), with 7 of 10 cSCI rats meeting the clinical criteria for orthostatic hypotension (71) (Fig. S3A; two-way repeated ANOVA, F[6, 102] = 9.10, *p* < 0.001; Fig. S3B. Two-way repeated ANOVA, F[6, 102] = 9.83, *p* < 0.001). There were no significant differences in pulse pressure (PP) across groups during either tilt position, in addition to no differences in HR, MAP, SBP, or DBP across groups during H-DT (we treated H-DT as a control change in position that did not generally challenge sympathetic outflow; Fig. 5C & 5D; Fig. S3C - S3F). Overall, these results indicate that cSCI impairs sympathoexcitatory capacity during orthostatic stress, leading to hypotension and impaired cardiovascular function.

**Fig. 5.**
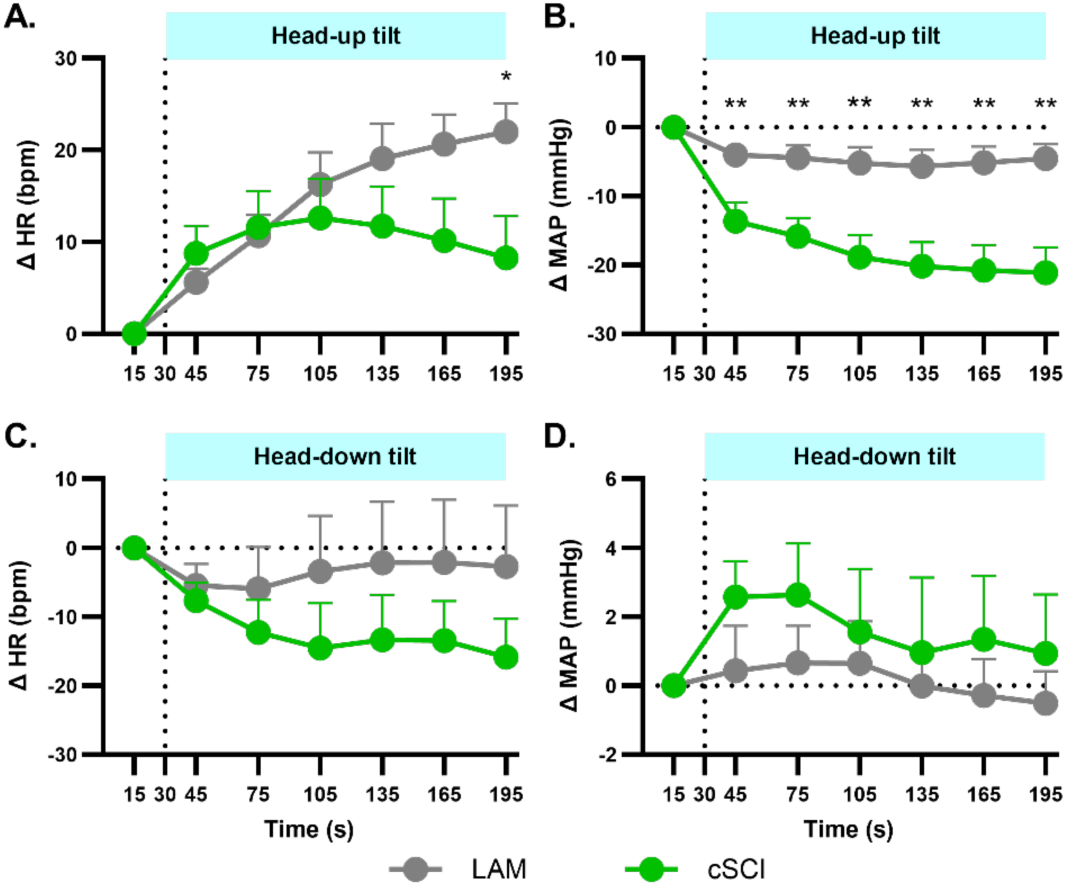
cSCI Significantly Disrupts the Control of Heart Rate and Mean Arterial Pressure During Orthostatic Stress. During orthostatic stress (i.e., head-up tilt), cSCI rats demonstrated significant decreases in heart rate (HR; **A**) and mean arterial pressure (MAP; **B**) compared to LAM rats, indicating a diminished sympathoexcitatory response due to spinal cord damage. As expected, there were no significant differences in HR (**C**) or MAP (**D**) during head-down tilt across groups (bpm: beats per minute; mmHg: millimeters of mercury; vertical dotted lines = time of given type of tilt; light blue shaded header = given type of tilt; \**p* < 0.05; \*\**p* < 0.01; mean ± S.E.M.).

**Fig. S3.**
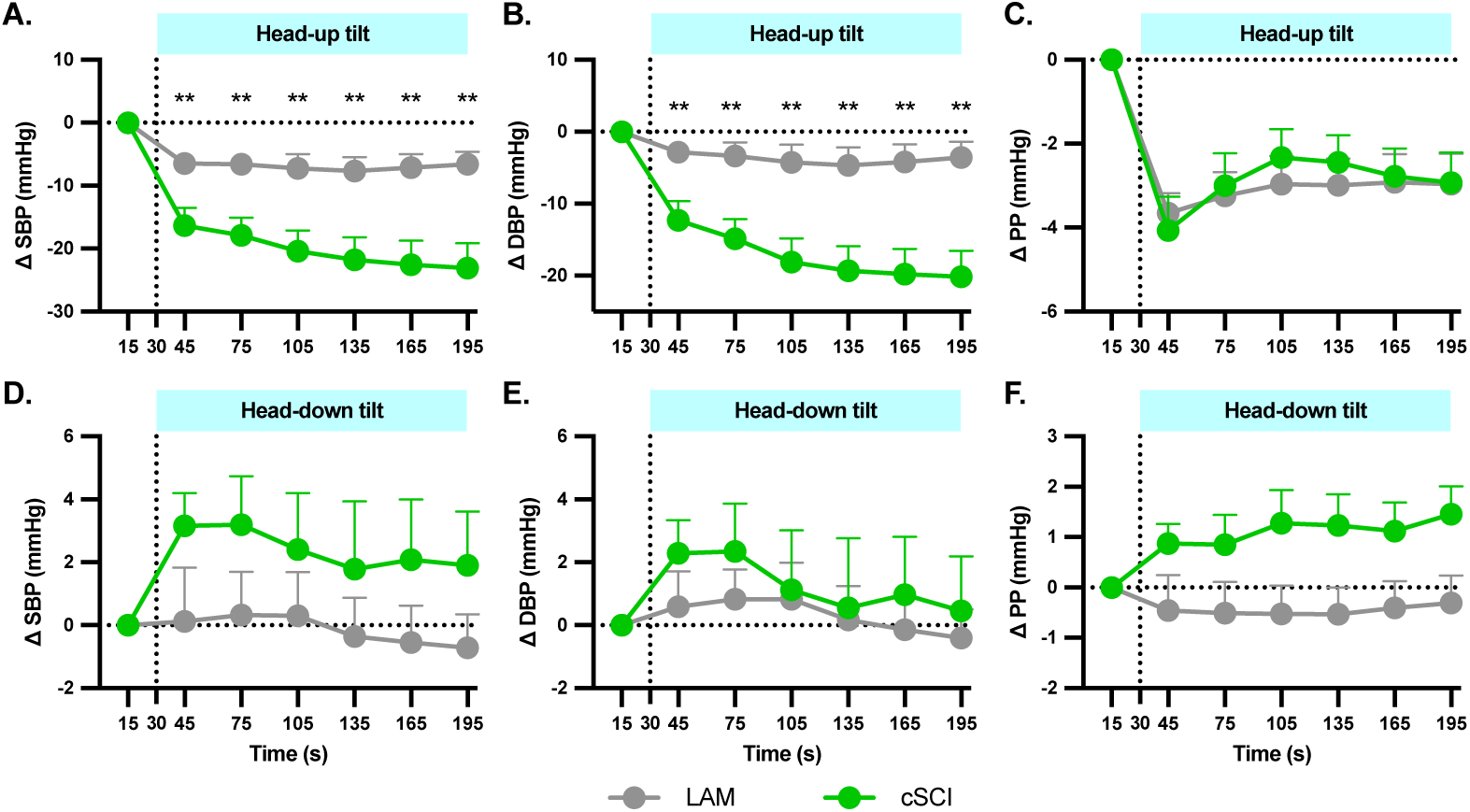
cSCI Significantly Impairs Blood Pressure During Systole and Diastole. During orthostatic stress (i.e., head-up tilt), cSCI rats demonstrated significantly lower systolic blood pressure (SBP; **A**) and diastolic blood pressure (DBP; **B**) compared to LAM rats, but not pulse pressure (PP; **C**). There were no significant differences across groups in SBP (**D**), DBP (**E**), or PP (**F**) during head-down tilt (mmHg: millimeters of mercury; vertical dotted lines = time of given type of tilt; light blue shaded header = given type of tilt; \*\**p* < 0.01; mean ± S.E.M.).

Impedance cardiography was also used to assess correlates of inotropy and stroke volume. During H-UT, C index, SV index and CO index demonstrated significant decreases in cSCI compared to LAM (Fig. 6A - 6C). Again, as expected, there were no changes in any metric during the H-DT control across groups (Fig. 6D – 6F). These results support the hypothesis that cSCI impairs CO index and its determinants. An overview of the orthostatic stress testing results is presented in Figure S5.

**Fig. 6.**
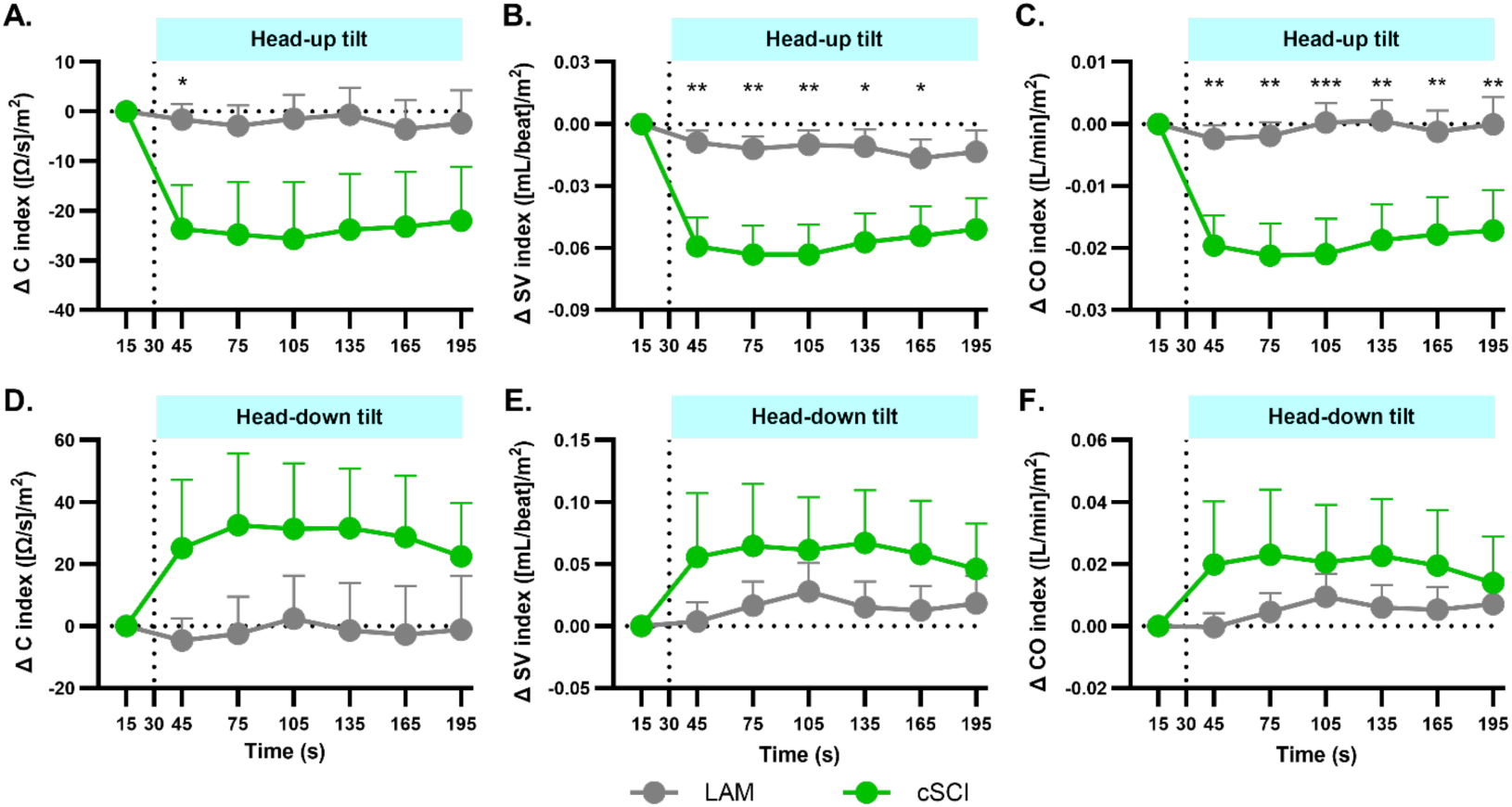
cSCI Significantly Disrupts Myocardial Contractility Index, Stroke Volume Index and Cardiac Output Index During Orthostatic Stress. During orthostatic stress (i.e., head-up tilt), cSCI rats showed significant decreases in myocardial contractility index (C index; **A**), stroke volume index (SV index; **B**) and cardiac output index (CO index; **C**), compared to LAM rats. There were no significant differences across groups for C index (**D**), SV index (**E**), or CO index (**F**) during head-down tilt. (Ω: ohms; vertical dotted lines = time of given type of tilt; light blue shaded header = given type of tilt; \**p* < 0.05; \*\**p* < 0.01; \*\*\**p* < 0.001; mean ± S.E.M.).

**Fig S5.**
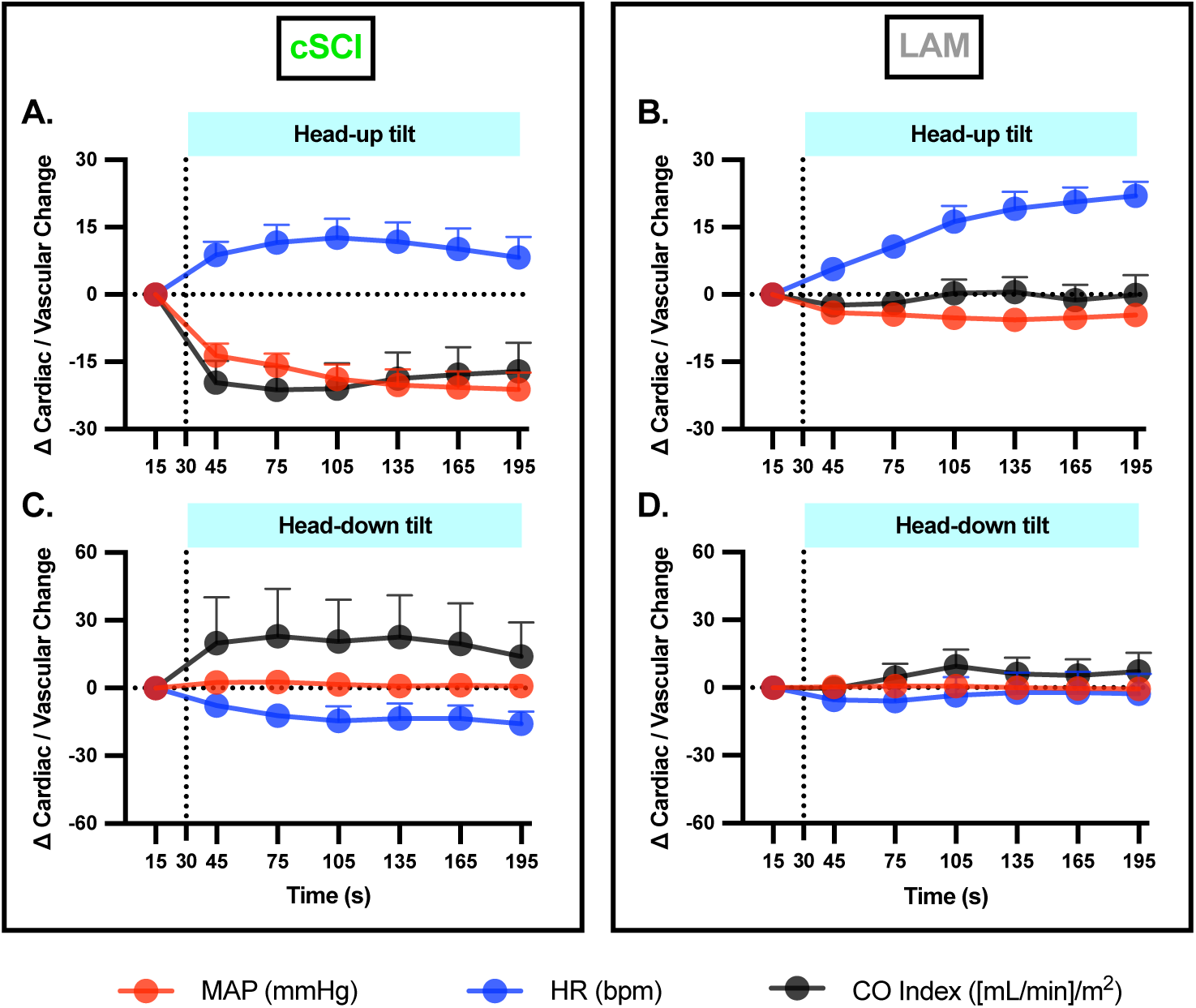
Orthostatic Stress Testing Results Overview (Δ Cardiac & Δ Vascular Metrics Superimposed Across Time). cSCI animals (**A**) exhibited significantly dysregulated cardiac & vascular responses compared to LAM (**B**) during H-UT. As expected, little to no changes occurred during H-DT across groups (**C** & **D**) (MAP: mean arterial pressure; HR: heart rate; CO index: cardiac output index; bpm: beats per minute; mmHg: millimeters of mercury; vertical dotted lines = time of given type of tilt; light blue shaded header = given type of tilt; mean ± S.E.M.).

### cSCI Impacts the Ability to Increase Stroke Volume Index and Cardiac Output Index During Pharmacologically Mediated Stress

Cardiovascular function was next assessed during pharmacological stress testing using infusions of dobutamine (DOB) (Fig. 1, bottom right), an agent commonly used to challenge the cardiovascular system (34,36–38). We assessed the hypothesis that cSCI would significantly impair the achieved SV index and CO index during DOB infusion. Fig. 7A, 7C, 7E & 7G show cardiovascular metric changes across time during pharmacological stress testing (HR, MAP, SV index, and CO index, respectively). Across groups, there were no significant differences in any cardiovascular metric during the low dose infusion of DOB (2 µg/kg/min; Fig. 7B, 7D, 7F, & 7H).

**Fig. 7.**
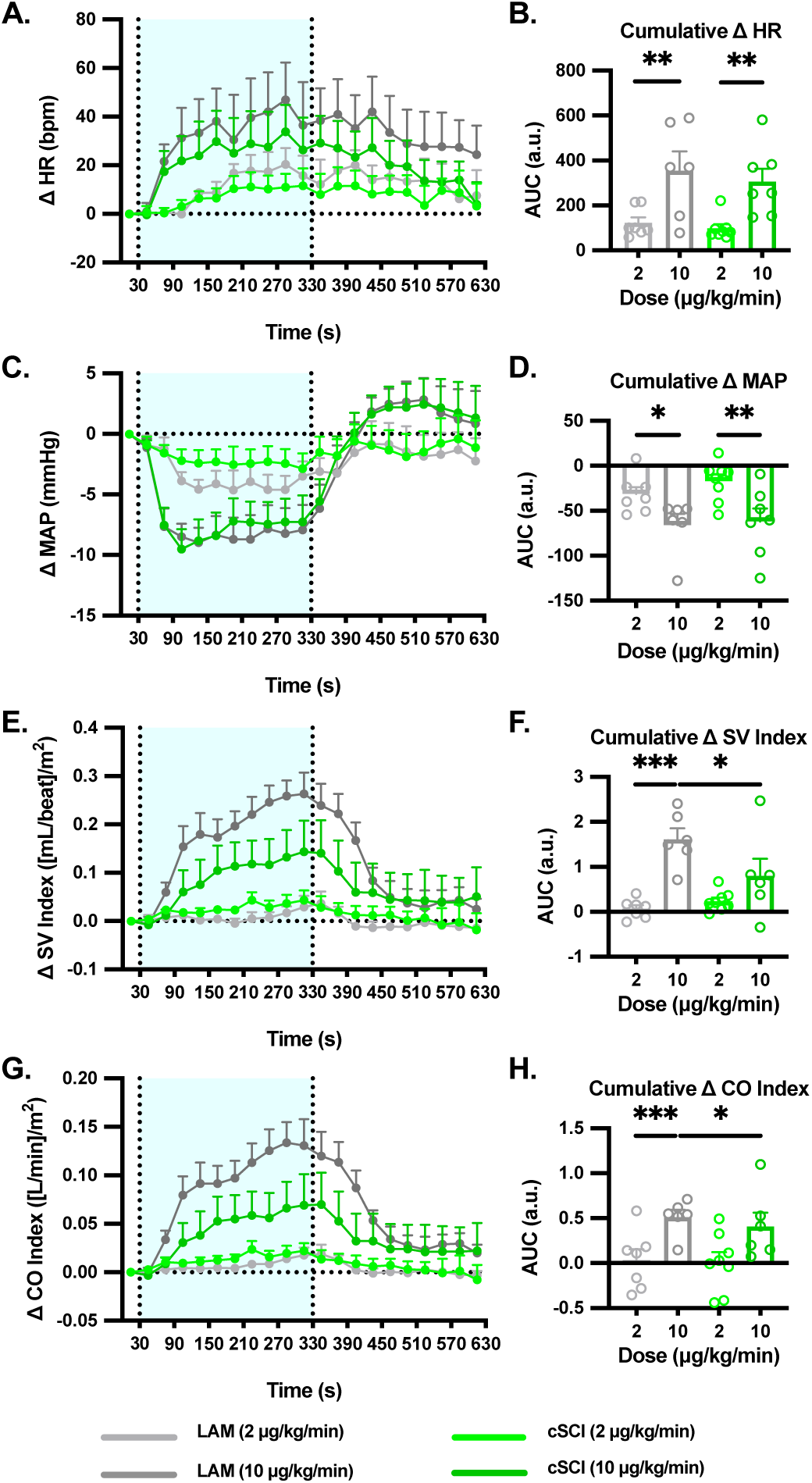
cSCI Signficantly Impairs the Ability to Increase Stroke Volume Index and Cardiac Output Index During Pharmacological Stress. During dobutamine (DOB) infusion, there were no significant differences in heart rate (HR; **A** and **B**) or mean arterial pressure (MAP; **C** and **D**) across groups. Importantly, cSCI rats achieved a significantly lower stroke volume index (SV index; **E** and **F**) and cardiac output index (CO index; **G** and **H**) compared to LAM rats at the higher DOB infusion dose level (AUC: area under the curve, a.u.: arbitrary unit, bpm: beats per minute, mmHg: millimeters of mercury; vertical dotted lines = time of given type of infusion; light blue shaded header = given type of infusion; \**p* < 0.05; \*\**p* < 0.01; \*\*\**p* < 0.0001; mean ± S.E.M.).

This is likely due to the milder pharmacological stress associated with this dose level. As expected, general dose dependent increases were observed within groups for HR, MAP, SV Index, and CO Index (Fig. 7B, 7D, 7F, and 7H). During the high dose infusion of DOB (10 µg/kg/min), cSCI rats achieved significantly diminished levels of SV index and CO index compared to LAM (Fig. 7F; two-way ANOVA, F[1, 23] = 4.66, *p* = 0.0420; Fig. 7H; two-way ANOVA, F[1, 23] = 5.56, *p* = 0.0270). Overall, these results provide further evidence of cardiovascular dysfunction, and importantly complement the impaired SV index and CO index results observed during orthostatic stress (using a different form of physiological challenge).

## Discussion

This study assessed the effects of chronic incomplete bilateral cSCI on aerobic function, muscle endurance, and cardiovascular control in rats. Using exercise stress testing, we observed significant effects of cSCI on aerobic fitness (Fig. 3) and forelimb muscle endurance (Fig. 4). In addition, there were numerous alternations to cardiovascular function following preclinical cSCI. cSCI rats had significantly reduced SBP at baseline (Table 1), similar to previous reports assessing cardiovascular function following SCI in humans or rats (8, 72–74). During orthostatic stress, the cardiovascular impairments were distributed impacting HR, MAP, C index, SV index, and CO index (Fig. 5 & Fig. 6). There was also significant cardiovascular dysfunction observed in cSCI rats during DOB infusion (Fig. 7), in part complimenting the deficits observed during orthostatic stress. Overall, these results provide extensive and clinically relevant evidence of the distributed impact of preclinical cSCI on multiple physiological systems that are critical for appropriately responding to physiological challenges. More importantly, these findings characterize in detail a preclinical model of SCI that exhibits clinically relevant deficits in aerobic, muscular, and cardiovascular function.

### cSCI’s Impact on Aerobic Fitness and Muscle Endurance

Peak oxygen consumption (VO_2 peak_) is a measure of aerobic capacity and overall cardiorespiratory fitness (20–23). Interestingly, cSCI rats only demonstrated a mild trend towards a decrease in VO_2 peak_ compared to LAM (Fig. 3A). Importantly, there are several physiological substrates contributing to VO_2_. The inspiratory muscles are mainly innervated by the phrenic motoneurons from C3 to C6 in rats (75–77). Previous studies demonstrate that a high-level SCI (e.g., a C2 hemicontusion) can disrupt diaphragm control. Regardless, largely preserved total ventilation is commonly observed in this model of SCI, potentially due to the amount of spared spinal tissue (75). A mid-cervical SCI study in rats also demonstrated that respiratory function progressively returns following injury (78), complimenting the findings of other similar studies (79,80). Therefore, respiratory capacity in our cSCI model may have been preserved (especially considering the extended 13 week recovery period), possibly contributing to the generally preserved VO_2 peak_ levels compared to LAM. Future studies will need to employ direct assessment of pulmonary and respiratory function to address these and other hypotheses.

The volume of aerobically active muscle during exercise stress testing is also a significant contributor to VO_2 peak_. cSCI rats exhibited BBB scores indicative of sufficient residual hindlimb function and consistent weight supported stepping (Fig. S1). Furthermore, cSCI and LAM rats achieved similar final running speeds during metabolic treadmill testing (18.4 ± 0.8 m/min vs. 20.3 ± 0.9 m/min, respectively). These results indicate that our model of cSCI critically resulted in enough residual locomotor capacity allowing for animals to complete the maximal aerobic exercise treadmill assessment, with the general trade-off of retaining a significant amount of aerobically active muscle. The preserved locomotor capacity of cSCI rats in this study is in part due to 1) white matter sparing in cSCI rats (Fig. 2C, right), and 2) the thoracolumbar locomotor central pattern generator (CPG) in this quadruped. The locomotor CPG in rats is distributed in the lower thoracic and lumbar spinal cord, and can generate rhythmic hindlimb movement, even in the presence of little to no supraspinal input for facilitating locomotion (81,82). The locomotor CPG is completely spared in our injury model, in addition to descending inputs sufficient for enabling consistent weight-supported stepping. Furthermore, we hypothesize that the active hindlimb muscles may have also facilitated increases in venous return due to the skeletal muscle pump, likely further contributing to the generally preserved VO_2 peak_ levels observed. Overall, future studies may assess modifying SCI severity, as well as integrating cardiovascular telemetry recordings during aerobic exercise testing, to further extend these findings.

Importantly, VO_2 peak_ was significantly negatively correlated with EPOC, in addition to a significant increase in EPOC observed in cSCI animals (Fig. 3B). Overall, these results indicate impaired aerobic fitness following cSCI. EPOC reflects the additional oxygen consumption needed during exercise recovery, with EPOC being lower when aerobic capacity and overall fitness is high (28,29; also see correlation analysis in results). EPOC specifically facilitates the replenishment of oxygen in the muscles and blood, in addition to phosphagen (ATP and creatine phosphate) restoration (24–29). The EPOC increase observed in cSCI animals may be due to multiple mechanisms, including disruption of sympathetically-mediated cardiovascular function impacting oxygen delivery, decreased mitochondrial density, or impaired thermoregulatory function leading to increased oxygen needs post-exercise (24,27,32,83–85).

Regarding resistance exercise stress testing, we used the isometric pull task to examine measures of forelimb fitness, including strength and muscle endurance. The vast majority of forelimb muscles engaged during this task are innervated by spinal levels above C8. Therefore, a C8 contusion should largely preserve the forelimb motor control circuits above the lesion (86). Overall, our results support this hypothesis, as forelimb strength was not impacted in cSCI animals (Fig. 4C). However, we did observe significant decreases in maximal force velocity following cSCI (Fig. 4B).

We also utilized the isometric pull task to examine muscle endurance, similar to previous muscle endurance studies using maximal voluntary contraction resistance exercise, requiring near maximal force exertion and high trial counts over time (44–47). Muscle endurance describes the capacity of a muscle or group of muscles to sustain repeated contractions against a load over an extended duration (87). Therefore, our muscle endurance measure is well aligned with this standard definition, as we quantified the change in peak force from the beginning to the end of the 30-minute testing session (involving an average of 515 ± 65 isometric pulls total during a given 30-minute testing session). cSCI rats demonstrated significantly reduced muscle endurance compared to LAM, with no change in the total number of trials across groups (Fig. 4). We hypothesize that this deficit in muscle endurance is in part accounted for by the cardiovascular deficits following cSCI (discussed in ‘*<u>cSCI Impacts Multiple Facets of Cardiovascular Function During Orthostatic Stress Testing</u>*’ below). Regardless, many other factors should be investigated to further elucidate the mechanisms of impaired forelimb muscle endurance following cSCI, including the accumulation of lactic acid, mitochondrial dysfunction, muscle fiber type switching, and oxygen delivery dysfunction (88).

### The Impact of cSCI on Cardiovascular Control Circuits

Cervical SCI often results in cardiovascular dysfunction due to the interruption of the descending sympathoexcitatory pathways. Sympathetic preganglionic neurons controlling cardiovascular functions are located in the intermediolateral (IML) columns spanning the thoracic (T1-T13) and upper lumbar spinal segments (L1-L2) in the rat (2,3). Sympathetic innervation of the heart and vasculature of the upper limbs generally originates from T1-T4, whereas the splanchnic vasculature and blood vessels of the lower limbs are generally innervated by spinal levels T6-L2. Therefore, a contusion at spinal level C8 significantly interrupts these descending sympathoexcitatory pathways regulating cardiac and distributed vascular function below the level of injury.

An extensive and excellent set of studies exist that assess the impacts of high thoracic SCI on cardiovascular control (10,72,73,89–94). The main differences between the current study and these additional studies are: 1) the level of the SCI (lower cervical vs. high thoracic) and 2) the severity of the SCI. Regardless, there should be only minor to modest differences using either lesion approach with respect to subsequent cardiovascular effects. For example, we observed a significant decrease in SBP and a trend towards a decrease in MAP at rest in cSCI rats (Table 1), similar to previous studies using a high thoracic SCI (72,73,89). We also demonstrate that cSCI leads to orthostatic hypotension and orthostatic dysregulation during baroreceptor unloading, similar to high thoracic SCI, albeit using different methods (please see ‘*<u>Comparing Head-Up Tilting, Lower Body Negative Pressure, and Other Methods Used to Assess Cardiovascular Responses to Baroreceptor Unloading’</u>*). Importantly, we observed a diminished ability to significantly increase HR during head-up tilting and baroreceptor unloading in cSCI rats (Fig. 5A), in contrast to other studies demonstrating a general persevered ability to increase heart rate following spinal lesion (72,73,89). In these additional studies, a portion of the T1-T4 cardiac IML column is still supraspinally innervated following an upper thoracic SCI. Furthermore, several studies also demonstrate an increased innervation to the IML following SCI within or below T1-T4, possibly due to sprouting of damaged sympathoexcitatory fibers at and above the level of the given lesion in part regulating cardiovascular function (89,95). Whether this spared innervation to the heart and other cardiovascular tissues significantly mediates these differential effects on heart rate control requires further investigation. Regardless, there are a variety of spinal lesion types that can result in clinically relevant cardiovascular dysfunction.

### cSCI Impacts Multiple Facets of Cardiovascular Function During Orthostatic Stress Testing

Our orthostatic stress testing results extend findings from numerous previous studies using head-up tilting (H-UT) in rats to induce significant hypotension (Table 2). Interestingly, several of these investigations demonstrate that significant hypotension can even be achieved during H-UT by simply using pharmacological agents that block the effects of sympathetic outflow (96–99). Overall, the MAP changes in these studies ranged from approximately −10 mmHg to −30 mmHg during various degrees of H-UT. Similar to our current study, significant hypotension can be achieved by damaging neural circuits that transmit sympathoexcitatory signals to the target cardiovascular tissues (72,75,89,100,101). The work by Lujan and DiCarlo 2020 is of particular importance (89), as it demonstrates that orthostatic dysregulation is present in the awake animal following different levels of SCI using H-UT (e.g., orthostatic dysregulation occurred following a cervical transection SCI, as well as a midthoracic transection SCI). Overall, the range of H-UT angles used in these studies spanned 45° – 90° (Table 2), similar to the ∼90° H-UT used in this study (for inducing a maximal orthostatic stress).

**Table 2.** Overview of published rat orthostatic stress studies.

| **Study Reference** | **Sex & Species** | **Anesthetized or Awake** | **aBP Measurement** | **Method for Inducing Orthostatic Dysregulation** | **HUT Angle** | **Approximate $\Delta$ mmHg** |
| --- | --- | --- | --- | --- | --- | --- |
| Lujan & DiCarlo, 2020 (89) | Male Sprague Dawley | Awake | Via left carotid artery (placed in descending aorta) | Complete spinal transection at C6/C7 | 90° | -30 mmHg (at 1 min post-H-UT; metric: MAP) |
| Inchan et al., 2022 (102) | Male Wistar | Anesthetized | Via left common carotid artery | Aged rats (~72 weeks old) | 90° | -15 mmHg (at 1 min post-H-UT; metric: MAP) |
| Sato et al., 2002 (103) | Male Sprague Dawley | Anesthetized | Via right femoral artery (placed in aortic arch) | Neural control disruption of the abdominal splanchnic vasculature | 90° | -52 mmHg (at 10 s post-H-UT; metric: SBP) |
| Akiyama et al., 2002 (96) | Male Sprague Dawley | Anesthetized | Via left common carotid artery (placed at level of heart) | Tamsulosin (10 µg/kg; i.v.) | 45° | -25-30 mmHg (at 2 minutes post-H-UT; metric: SBP) |
| Akiyama et al., 2002 (96) | Male Sprague Dawley | Anesthetized | Via left common carotid artery (placed at level of heart) | Prazosin (10 µg/kg; i.v.) | 45° | -25-30 mmHg (at 2 minutes post-H-UT; metric: SBP) |
| Matos de Mora et al., 2005 (97) | Male Wistar | Anesthetized | Via left femoral artery (placed at level of heart) | Losartan (1 mg/kg; i.v.) | 90° | -35% (at 2 minutes post-H-UT; metric: % change in MAP) |
| Bedette et al., 2008 (98) | Male Wistar | Awake | Via femoral artery (placed at level of heart) | Telmisartan (10 mg; oral) | 75° | -10 mmHg (at 7.5 minutes post-H-UT; metric: MAP) |
| Funakami et al., 2010 (104) | Male Wistar | Anesthetized | Via left common carotid artery | Specific alteration of rhythm in temperature stress | 60° | -20 mmHg (at 2 minutes post-H-UT; metric: MAP) |
| Funakami et al., 2007 (99) | Male Wistar | Anesthetized | Via left common carotid artery | Hexamethonium (5 mg/kg; i.v.) | 60° | -17 mmHg (at 3 minutes post-H-UT; metric: MAP) |
| Nishimure et al., 2018 (105) | Male Sprague-Dawley | Anesthetized | Via right common carotid artery | Sinoaortic denervation | 90° | -12 mmHg (at 30 minutes post-H-UT; metric: MAP) |
| Karakida et al., 1991 (106) | Male Wistar | Anesthetized | Via caudal artery in the tail | Streptozotocin diabetes | 45° | -24 mmHg (at 20~25 seconds post-H-UT; metric: SBP) |
| Lee et al., 1982 (107) | Male Sprague-Dawley | Awake | Via femoral artery (placed at level of heart) | Guanethidine (0.03 mg/kg; i.v.) | 90° | -18 mmHg (at 2 minutes post-H-UT; metric: MAP) |

Following cSCI in the current study, positional change from supine to H-UT resulted in significant hypotension, with 7 of 10 rats meeting the criteria for clinical orthostatic hypotension (71) (Fig. 5B). Numerous studies demonstrate that in uninjured subjects, descending sympathetic outflow controlling the heart and vasculature compensates for blood pooling in the lower body and diminished cerebral perfusion during H-UT, allowing for near normal blood pressure and tissue perfusion (2,108–110). Unfortunately, following cSCI communication between the descending sympathetic outflow pathways and target tissues is significantly disrupted, leading to diminished control of the heart and vasculature. In addition to vascular contributions, it is likely that orthostatic dysregulation also occurred in part due to mildly impaired cardiac function. During H-UT, cSCI animals demonstrated an inability to increase in HR, along with significant decreases in C index, SV index, and CO index compared to LAM (Fig. 5A, 6A, 6B, & 6C). Overall, the differential and complex contributions of cardiac (e.g., left ventricular atrophy following SCI (111–113)) and hemodynamic factors (e.g., venous pooling diminishing preload and SV following SCI (114–116)) require further investigation.

Despite the partial loss of descending sympathetic outflow to the heart, HR increased initially, followed by an overall inability to maintain this increase by the end of the H-UT period (Fig. 5A). We hypothesize that the initial increase in HR is likely accounted for by baroreceptor-mediated modulation via spared pathways as well as vagal withdrawal. Baroreceptors are a type of mechanoreceptor sensitive to changes in aBP, located in tissues such as the carotid sinus and aortic arch (12). An increase or decrease in aBP will activate the baroreflex, with subsequent inhibition or enhancement of sympathetic outflow, respectively (12). The significant decrease in MAP during H-UT (Fig. 5B) may activate the baroreflex, increasing sympathoexcitation via residual descending sympathoexcitatory pathways partially elevating HR. In addition, H-UT likely also leads to reflexive vagal withdrawal, facilitating partial increases in HR.

No groups demonstrated cardiovascular deficits during the control position of H-DT (Fig. 5C, 5D, & 6D-6F). This was not unexpected, as H-DT partially mimics the Trendelenburg position that may have minimal effects in general on the overall cardiovascular parameters measured in this study (117). Furthermore, H-DT may primarily engage the parasympathetic system which is spared following cSCI.

### Impedance Cardiography for Estimating Stroke Volume Index in Rats

Impedance cardiography (ZCG) measures the thoracic impedance associated with the cardiac cycle. The derivative of the ZCG signal 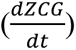 has been used in numerous studies to derive hemodynamic metrics such as SV and CO (51–53,118). The ZCG-derived SV was adjusted by body surface area to account for different sizes of animals. Since ZCG has been introduced to monitor hemodynamics, most of its applications have focused on human subjects (51,55,118–120), in addition to dogs (54,55,121–123) and pigs (124–126). Our current study extends previous studies using ZCG in rats (49,50,56). Due to the rapid cardiac cycle in rats, ZCG may miss valuable hemodynamic features, such as aortic valve opening and closing (51) that are in part used to estimate LVET. This shortcoming was overcome by simultaneously recording the aBP waveform, providing a gold standard means for quantifying LVET (127–129) (Fig. 1, bottom middle). IN addition, despite the acceptable hemodynamic parameter estimation accuracy that ZCG provides, it is still important to validate its derived measures using other well-accepted techniques. ZCG has been widely validated compared to a number of other techniques for measuring SV and other parameters (51,55,57,130). Finally, there are different formulas for calculating SV and SV index from ZCG, and the definition of each variable may slightly differ across studies (49,51–53,118,130). Our study calculated 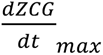 using the difference in impedance derivative from the beginning of systole to the signal peak, divided by the time interval, similar to previous works (56,58,59). Other formulas may use the maximal rate of impedance change to determine 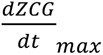. Future studies are needed to further validate hemodynamic measurements derived from ZCG in rats.

### cSCI Impairs Cardiovascular Responses During Pharmacological Stress

Dobutamine (DOB) is a well-studied β1 and β2 adrenergic receptor agonist. Via β1 receptor activation, it has a strong inotropic effects for increasing SV and a relatively modest chronotropic effect for increasing HR (131,132). In addition, DOB exhibits vasodilatory effects via β2 receptors. The use of DOB has been used in many preclinical studies to evaluate cardiocascular function (34,36–38) and is also extensively used as a pharmacological stressor during echocardiography recordings (34,35,133,134). Therefore, DOB was used in this study to examine cardiovascular capacity following cSCI, similar to our previous study in uninjured rats (38). We hypothesized that pharmacological stress using DOB infusion would reveal cardiovascular dysfunction in cSCI rats. As expected, HR, MAP, SV index, and CO index showed significant dose-dependent changes in both cSCI and LAM rats (Fig. 7). Importantly, cSCI rats achieved a significantly smaller SV index and CO index during high dose DOB infusion compared to LAM rats (Fig. 7F & 7H). These impairments may in part be attributed to a lower baseline NE plasma concentration (9,135), in addition to as left ventricular atrophy and general cardiac remodeling following SCI (111–113). Overall, there were larger impairments in SV index and CO index during DOB stress testing compared to orthostatic stress testing. Regardless, these pharmacological stress testing results provide independent verification of impaired cardiovascular capacity following cSCI, complimenting the orthostatic stress testing results.

### Comparing Head-Up Tilting, Lower Body Negative Pressure, and Other Methods Used to Assess Cardiovascular Responses to Baroreceptor Unloading

Multiple techniques have been developed to modulate arterial baroreceptor loading and evaluate cardiovascular control, such as lower-body negative pressure, neck suction, the modified Oxford technique, and the tilt table technique used for orthostatic stress testing in this study. Lower-body negative pressure applies subatmospheric pressure to the lower limbs sealed in an air tight chamber, reducing central venous pressure and venous return, mimicking baroreceptor unloading seen during orthostatic stress (72). Neck suction is another technique and uses pneumatic pressure applied to the cervical region, mechanically modulating the carotid baroreceptors and triggering baroreflex cascades (136). The modified Oxford technique provides a pharmaceutical approach to affect baroreflex function via separate intravenous infusions of a vasodilator (e.g., sodium nitroprusside) and a vasoconstrictor (e.g., phenylephrine) (137). Orthostatic stress testing does not use artificial or pharmaceutical manipulation to trigger the baroreflex, but instead uses natural changes in body position and gravity induced blood pooling in the lower limbs. Head-up orthostatic stress testing can also use different inclination angles, allowing for controlled transitions from supine to an upright position (Table 2).

Numerous studies demonstrate that orthostatic testing can be used to induce orthostatic stress and trigger cardiovascular dysregulation rats (Table 2). Regardless of the technique used, the hydrostatic indifference point must be considered to ensure optimal placement of the given arterial blood pressure catheter (138–141). Many studies have demonstrated that recording aBP from the hydrostatic indifference point (commonly located below or at the level of the heart) can interestingly, and dramatically, blunt blood pressure changes induced by changes in body position (138,141,142). Our blood pressure catheter was placed in the aortic arch above the heart, where numerous pressure sensitive aortic baroreceptors are located. Future studies should consider the potential effects of blood pressure catheter location within the vascular network on aBP measures during changes in body position and / or baroreceptor manipulation.

### Conclusions

Overall, these results describe a clinically relevant model of distributed physiological dysfunction following cervical SCI. Our experiments utilized an array of complimentary stress tests to characterize the effects of cervical SCI on both correlates of exercise tolerance and cardiovascular control. Our results are both internally consistent with regards to observed effects for a given metric, persisting across different kinds of stress testing (e.g., SV index and CO index impairments present during both orthostatic and pharmacological stress testing), as well as externally consistent when comparing our outcomes to other important preclinical (72,89,143) and clinical studies (6). From a methodological perspective, we implemented multiple levels of controls in our experiments (e.g., cSCI vs. LAM, H-UT vs. H-DT, or low vs. high dose DOB), in addition to robust measurement techniques (e.g., muscle endurance testing with high trial counts and gold standard aBP recorded from within the aortic arch). Future studies should assess the persistence of these findings in conscious rats (e.g., using radiotelemetry). In addition, the further assessment of histopathology following cervical SCI is warranted to investigate substrates of dysfunction at the tissue level.

## Acknowledgements

This study was supported by the National Institutes of Health under grant number 1R01NS131493.

